# Dimensionality changes actin network through lamin A and C and zyxin

**DOI:** 10.1101/752691

**Authors:** Jip Zonderland, Ivan Lorenzo Moldero, Carlos Mota, Lorenzo Moroni

## Abstract

The actin cytoskeleton plays a key role in differentiation of human mesenchymal stromal cells (hMSCs), but its regulation in 3D tissue engineered scaffolds remains poorly studied. hMSCs cultured on 3D electrospun scaffolds made of a stiff material do not form actin stress fibers, contrary to hMSCs on 2D films of the same material. On 3D electrospun- and 3D additive manufactured scaffolds, hMSCs also displayed fewer focal adhesions, lower lamin A and C expression and less YAP1 nuclear localization. Together, this shows that dimensionality prevents the build-up of cellular tension, even on stiff materials. Knock down of either lamin A and C or zyxin resulted in fewer stress fibers in the cell center. Zyxin knock down reduced lamin A and C expression, but not vice versa, showing that this signal chain starts from the outside of the cell. Our study demonstrates that dimensionality changes the actin cytoskeleton through lamin A and C and zyxin, an important insight for future scaffold design, as the actin network, focal adhesions and nuclear stiffness are all critical for hMSC differentiation.

## 1. Introduction

Understanding cellular responses to material properties (such as stiffness, chemistry, and topography) are critical endeavors in fields such as tissue engineering and regenerative medicine. Unraveling the molecular mechanisms underlying these cellular responses can lead to more intelligent design of tissue engineering constructs. ^[1]^ Cells adhere to materials through integrins, forming focal adhesion complexes that connect the actin cytoskeleton to the extra cellular matrix (ECM). ^[2]^ The actin filaments attach to other focal adhesions, or to the nucleus through the linker of nucleoskeleton and cytoskeleton (LINC) complex and lamin A and C. ^[3]^ Lamin A and C form a protein meshwork under the nuclear membrane to give structural integrity to the nucleus. ^[4–6]^ Actin filaments join together to form stress fibers, with incorporated non-muscle myosin to create contractile force between its two attachment points. ^[7]^ On stiffer materials, focal adhesions and stress fibers have been shown to be bigger and more abundant than on softer materials ^[8–11]^, creating a higher cellular tension. ^[11, 12]^ Indeed, lamin A and C expression, indirectly attached to actin stress fibers, has also been shown to increase on stiffer materials. ^[6]^ Besides material stiffness, other factors such as material chemistry and topography have also been shown to influence focal adhesions, actin stress fibers and lamin A and C. ^[13–15]^ Yes-associated protein 1 (YAP1) is an important mechanosensitive co-transcription factor that translocates to the nucleus at higher cellular tension to transduce these mechanical changes in the cell to changes in gene expression. ^[16]^ On stiffer materials and with more cellular tension human mesenchymal stromal cells (hMSCs) show increased osteogenic differentiation, while softer materials and lower cellular tension enhance differentiation to chondro- and adipogenic lineages. ^[17–21]^ These changes in hMSC differentiation have been shown to be orchestrated by actin stress fibers ^[22]^, focal adhesions ^[22]^, lamin A and C ^[23]^ and Yes-associated protein 1 (YAP1). ^[24, 25]^

While other material properties are relatively well studied, the dimensionality (2D v. 3D) of a material has not yet been widely studied. All 3D tissue engineering constructs inherently introduce dimensionality. It is therefore important to understand the effect of dimensionality on the proteins involved in cellular tension, as they are critical for hMSC differentiation. ^[22–25]^ Some studies have investigated the role of the dimensionality on cellular tension in soft hydrogels. ^[26]^ While this gives valuable insights for some tissue engineering applications, many 3D tissue engineering constructs are made of stiff materials. Therefore, we investigated the effect of dimensionality in 3D electrospun (ESP) and 3D additive manufactured (AM) scaffolds and compared to flat films made of the same stiff material (300PEOT55PBT45, PolyActive™, 100 MPa ^[27]^) (Figure 1a). Specifically, we focused on four main parts of the cellular tension machinery: focal adhesions (zyxin and paxillin), actin cytoskeleton, nuclear skeleton (lamin A and C) and a mechanosensitive co-transcription factor (YAP1).

**Figure 1.**
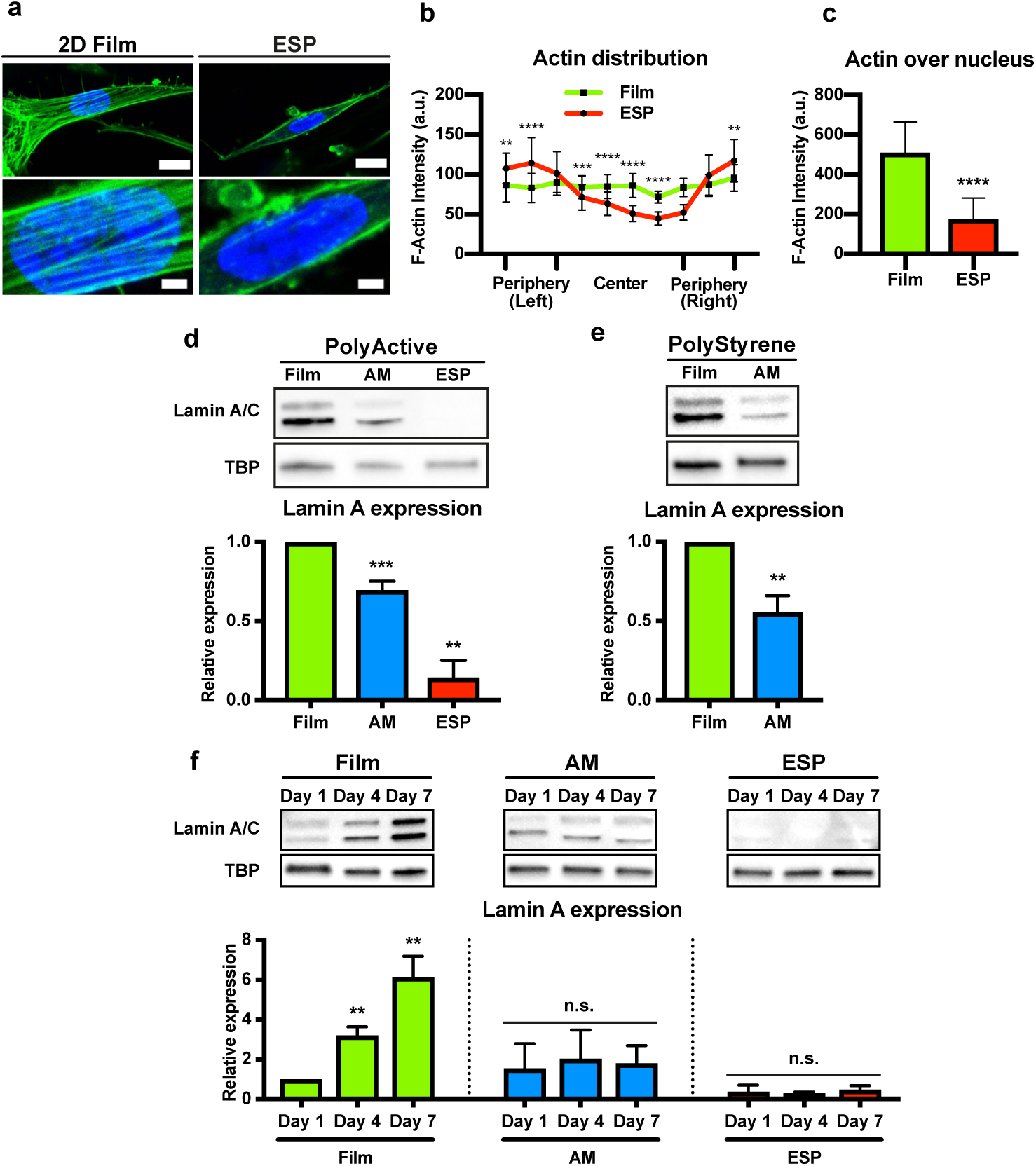
Different actin network organization and lower nuclear stiffness in hMSCs cultured on 3D ESP scaffolds. **a,** hMSCs cultured on 300PEOT55PBT45 (PolyActive™) 2D films and 3D electrospun (ESP) scaffolds stained for F-actin (green) and nuclei (blue). The bottom panels are magnifications (4×) of the respective images above. Scale bars represent 15 µm (top panels) and 3 µm (bottom panels). **b,** Quantification of the F-actin intensity distribution in hMSCs cultured on 2D films or 3D ESP scaffolds. n=21 for films and n=23 for ESP from 3 biological replicates. Two-way ANOVA; **p<0.01, ***p<0.001, ****p<0.0001. **c,** Quantification of F-actin intensity that overlaps with the nucleus in hMSCs cultured on 2D films or 3D ESP scaffolds. n=12 for films and n=13 for ESP from 3 biological replicates. Student’s t-test, ****p<0.0001. **d,** Lamin A expression in hMSCs grown on 2D films, 3D AM, and ESP scaffolds. TBP shown as loading control in the blots. Graph depicts average expression of lamin A/TBP normalized to films, quantified by western blots from 4 independent experiments. One-way ANOVA; **p<0.01, ***p<0.001 compared with films. **e,** Lamin A expression in hMSCs cultured on polystyrene, as 2D films or 3D AM scaffolds. TBP shown as loading control in the blots. Graph shows the average expression of lamin A/TBP normalized to films, quantified by western blot from 3 independent experiments. Ratio paired t-test; **p<0.01. **f,** Lamin A and C expression on day 1, 4 and 7 after seeding hMSCs on 2D films, 3D additive manufactured (AM) or electrospun (ESP) scaffolds. TBP shown as loading controls. Quantification of western blots from 3 independent experiments shows average expression of lamin A/TBP normalized to 2D films at day 1. One-way ANOVA; *p<0.05 compared with expression on 2D film at day 1.

We demonstrate that dimensionality influenced the actin stress fiber formation, focal adhesion formation, lamin A and C expression and YAP1 nuclear localization in these commonly used 3D tissue engineering scaffolds. The 3D scaffolds prevented the build-up of cellular tension, even on stiff materials, through a decrease in zyxin expression, which decreases lamin A and C expression and together change the actin cytoskeleton.

## 2. Results

### 2.1. Dimensionality prevents formation of actin stress fibers and reduces lamin A and C

To start investigating how dimensionality influences cellular tension in stiff materials, we looked at the actin cytoskeleton of hMSCs cultured on 3D ESP scaffolds or 2D films for 7 days (**Figure 1a**). As expected, on 2D films large actin stress fibers formed and F-actin fibers were distributed throughout the whole cell (Figure 1b). When hMSCs were cultured on ESP 3D scaffolds, we observed far fewer actin stress fibers and a change in F-actin distribution. F-actin fibers were mainly located on the cell periphery and very little F-actin was observed in the cell center or overlapping with the nucleus (Figure 1c). The lack of many stress fibers and a more pronounced peripheral actin network is a sign of lower cellular tension ^[28]^, even though cells were cultured on stiff materials. F-actin distribution of hMSCs in AM 3D scaffolds was attempted, but could not be evaluated due to the very high cell density in these 3D constructs. In this high cell density environment, the F-actin distribution of individual cells could not be discriminated from F-actin of surrounding cells.

To understand why few actin fibers were found overlapping with the cell nucleus, we evaluated the expression of lamin A and C, a protein that assists in linking actin filaments to the nucleus. ^[3]^ After 7 days of culture, lamin A and C expression were significantly reduced in AM scaffolds, by 30 ± 3.5% (p<0.001) and 46 ± 14% (p<0.05), respectively, compared to 2D films (Figure 1d and Figure S1a). Lamin A and C expression was even more reduced in hMSCs cultured on the ESP scaffolds, by 85 ± 6.5% (p<0.01) and 91 ± 8.0% (p<0.05), respectively, compared to those cultured on 2D films. The same trend was observed for polystyrene, a stiffer material: cells expressed 44 ± 4.7% (p<0.01) less lamin A and 71 ± 6.5% (p<0.01) less lamin C when cultured on AM 3D scaffolds, compared to 2D films (Figure 1e and Figure S1b).

To determine how lamin A and C expression changed over the culture period, we measured their expression levels at days 1, 4, and 7 (Figure 1f and Figure S2). Lamin A and C expression in hMSCs cultured on the 3D ESP or AM scaffolds remained low at all time points. In comparison, lamin A and C steadily increased over time in hMSCs cultured on 2D films. Lamin A was 3.2 ± 0.43 and 6.2 ± 1.0 times higher after day 4 and day 7, respectively, compared to day 1. Lamin C was 3.7 ± 1.0 and 8.4 ± 3.5 times higher on day 4 and day 7, respectively, compared to day 1. These findings show that due to the dimensionality, lamin A and C expression remains low over the culture period, in contrast to 2D, where lamin A and C expression increases over time.

### 2.2. Reduced nuclear stiffness of cells cultured in 3D

To test whether the change in lamin A and C expression affected nuclear stiffness, we observed hMSC migration from 2D films or 3D scaffolds, through a transwell with 3-µm or 8-µm pores. It has been shown that cells with high levels of lamin A and C (conferring greater nuclear stiffness) can migrate through 8-µm pores but are unable to move through 3-µm pores.^[29]^ Indeed, cells cultured on 2D films and harboring higher lamin A and C expression migrated less well through 3-µm pores than through 8-µm pores (p<0.05) (Figure 2a). Cells migrating from ESP scaffolds migrated equally well through 3-µm or 8-µm pores. The ratio of hMSCs migrating through 3-µm pores to those not migrating was significantly higher from ESP scaffolds than 2D films (p<0.01). hMSCs on AM scaffolds did not have sufficient physical contact with the transwells to migrate from the AM scaffold to the transwell membrane (data not shown), so their migration ability could not be evaluated. These data indicate that lower lamin A and C expression from hMSCS in 3D ESP resulted in lower nuclear stiffness of the cells, compared to hMSCs grown on 2D films.

**Figure 2.**
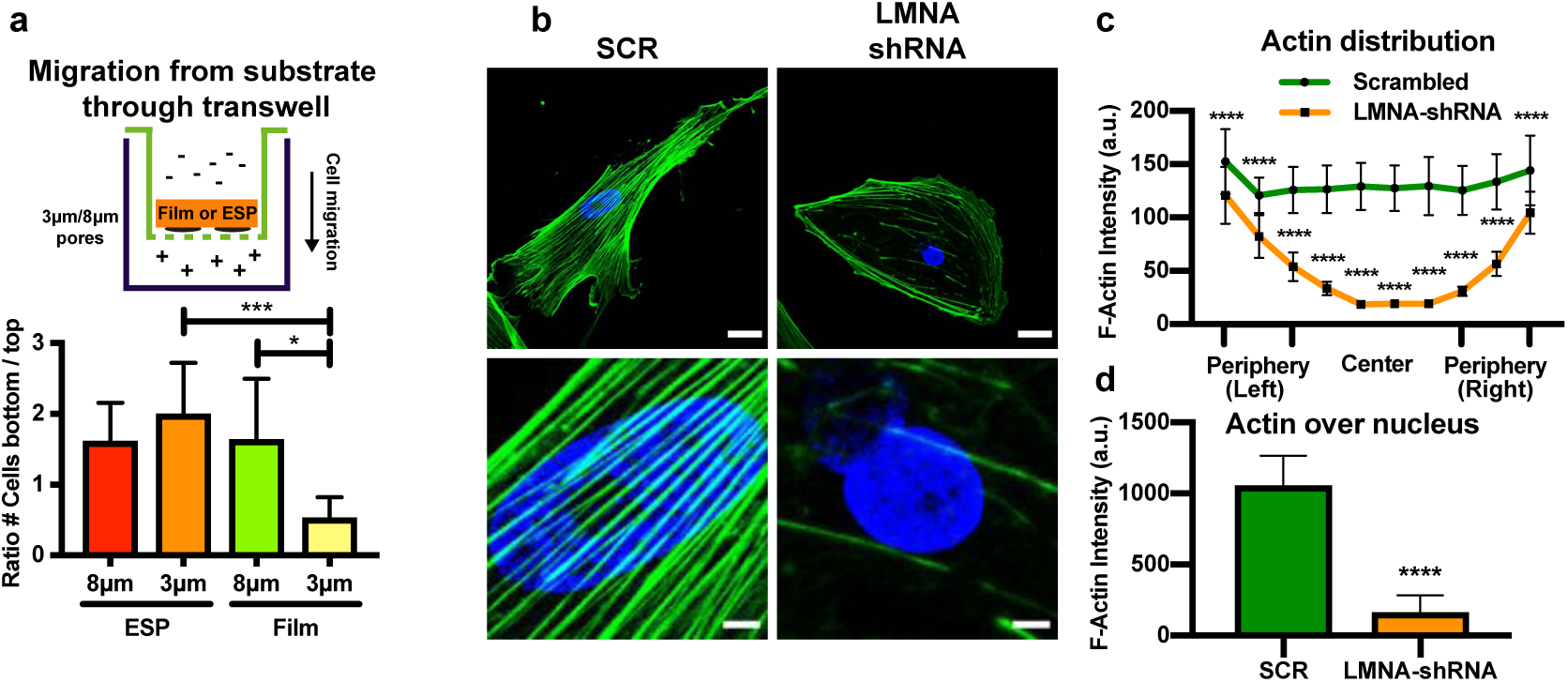
Lower nuclear stiffness in 3D ESP and Lamin A and C influences F-actin organization. **a,** Migration of hMSCs from 2D films or 3D ESP scaffolds through a transwell with 8-µm or 3-µm pores. 8 µm ESP: n=18; 3 µm ESP n=19; 8 µm Film n=18; 3 µm Film n=19, from 4 independent experiments.Krusal Wallis test; *p<0.05, ***p<0.001. For all graphs, error bars represent mean ± 95% CI. **b,** hMSCs transduced with scrambled-(SCR, left) or LMNA-shRNA (right), stained for F-actin (green) and nuclei (blue). The bottom panels are 7× magnifications of the respective images above. Scale bars represent 30 µm (top panels) and 5 µm (bottom panels). **c,** Quantification of the F-actin intensity distribution in hMSCs transduced with scrambled- or LMNA-shRNA. n=33 for SCR and n=29 for LMNA-shRNA from 3 biological replicates. Two-way ANOVA; ****p<0.0001. **d,** Quantification of F-actin intensity that overlaps with nuclei staining in hMSCs transduced with scrambled- or LMNA-shRNA. n=25 cells analyzed for SCR and n=22 for LMNA from 3 biological replicates. Mann-Whitney test, ****p<0.0001. Error bars represent mean ± SD (**b**) or mean ± 95% CI (**c, d**).

In addition to migration ability, nuclear stiffness was also assessed morphologically. While a stiff nucleus shows a uniform shape, lower nuclear stiffness results in more heterogeneous shapes and more ‘folds’. ^[30, 31]^ Indeed, nuclei of cells cultured in 2D showed mostly uniform nuclei, with folds in 33 ± 11% nuclei. From hMSCs on the 3D scaffolds, 76 ± 12% (p<0.01) and 96 ± 4% (p<0.0001) of nuclei displayed folds from AM and ESP scaffolds, respectively (Figure S3).

Together, these observations demonstrate that lower lamin A and C expression, resulting from the dimensionality of the 3D scaffolds compared to 2D films composed of the same stiff material, reduced nuclear stiffness, an observation not previously reported.

### 2.3. Lamin A and C plays a role in shaping the F-actin network

Actin tension has been shown to influence lamin A and C expression ^[32]^, but the role of lamin A and C in shaping the actin network is not known. To investigate this, cells were transduced with LMNA shRNA, knocking down both lamin A and C (Figure S4), and stained for F-actin (Figure 2b). A large change in actin organization was observed in the LMNA knock down. Similar to the actin organization of hMSCs on 3D ESP scaffolds, the LMNA knock down showed a clear decrease in F-actin in the cell center and over the nucleus (Figure 2c and d). Thus, lamin A and C plays an important role in shaping the actin network and could be partly responsible for the difference in actin organization in 2D vs 3D.

### 2.4. Cell density does not explain differences in lamin A and C expression observed between 2D and 3D

Because cell density inherently differs between 2D and 3D substrates, we wished to determine whether it was the reason for the differences we observed in lamin A and C expression. To determine the influence of cell density on lamin A and C expression, we cultured hMSCs for 7 days with different cell seeding densities on 2D TCP (**Figure 3a**). The different cell densities were chosen to reach confluency after 1 day (20k cells/cm^2^), 2 days (10k cells/cm^2^), or roughly 40%, 60%, 70%, or 80% confluency after 7 days (1.25k, 2.5k and 5k cells/cm^2^, respectively), or little cell contact after 7 days (625 cells/cm^2^). Increasing cell seeding density increased lamin A and C expression after 7 days of culture (Figure 3a and Figure S5a). At 20k cells/cm^2^, lamin A and C expression were 3.7 ± 1.7 (p<0.05) and 5.3 ± 1.7 (p<0.01) times higher than at 1.25k cells/cm^2^, respectively.

**Figure 3.**
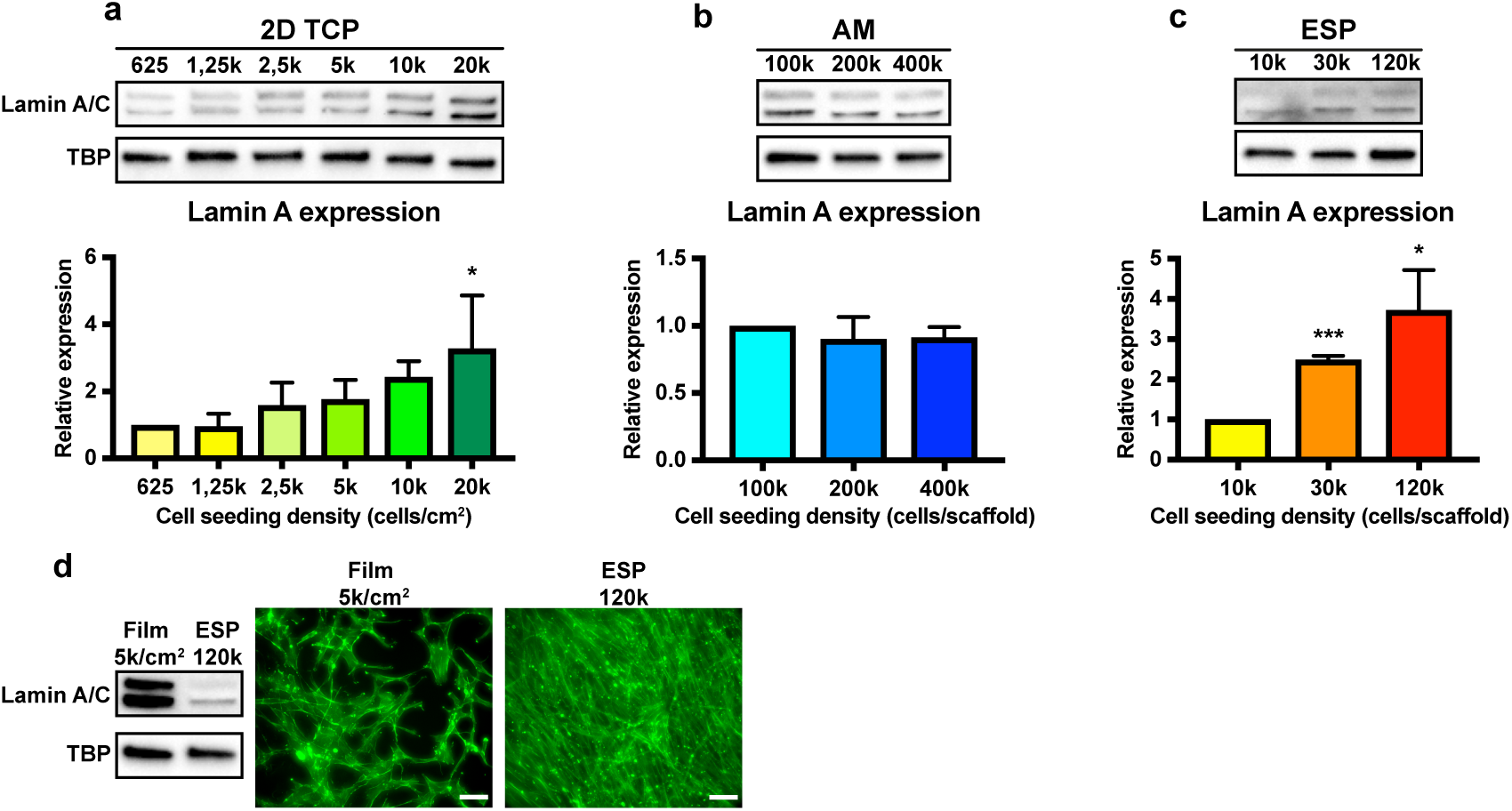
Cell seeding density influences lamin A and C expression. **a–c,** Lamin A and C expression of hMSCs seeded on 2D TCP (a), 3D AM (b), or 3D ESP (c) in varying densities. Western blots (top) and quantification of western blots (bottom) from 3 independent experiments. Error bars represent mean ± SD. Values were normalized to 625 cells/cm^2^ (a), 100k cells/scaffold (b), or 10K cells/scaffold (c). One-way ANOVA; *p<0.05 compared with 1250 cells/cm^2^ (a) or 10k cells/scaffold (c);. ***p<0.001, compared to 10k cells/scaffold. TBP shown as a loading control. **d,** Representative western blot (left) of hMSCs cultured on films at a medium cell density, or on electrospun scaffolds at a high cell density. Representative images of hMSCs on 2D films seeded at 5k cells/cm^2^ (left) and on a 3D ESP scaffold seeded with 120k cells (right) show the observed cell density at the time of harvest (day 7), visualized by F-actin staining (green). Scale bars represent 50 µm.

As lamin A and C were influenced by cell density in 2D, we next examined whether this was also true in the 3D scaffolds. For the 3D ESP scaffolds, different cell densities were chosen to have confluency after 1 day (120k cells/scaffold), near confluency after 7 days (30k cells/scaffold), or little cell contact after 7 days (10k cells/scaffold). For 3D AM scaffolds, confluency is more difficult to determine. With 400k cells/scaffold, most pores are filled after 1-2 days of culture. With 200k and 100k cells/scaffold, full pores are attained after 4–5 and 7 days of culture, respectively. Importantly, little cell proliferation was observed in 3D AM scaffolds (data not shown), thus the increased filling of the scaffold is not related to proliferation, unlike cells on 3D ESP or 2D TCP.

Similar to 2D TCP, increasing cell seeding density significantly increased lamin A expression in 3D ESP scaffolds, with 2.5 ± 0.1 (p<0.001) and 3.7 ± 1.0 (p<0.05) times more lamin A from 30k and 120k cells/scaffold, respectively, compared to 10k cells/scaffold (Figure 3c). Lamin C expression was not significantly changed by cell density on the ESP 3D scaffold, with 2.1 ± 1.0 and 1.9 ± 0.87 times more lamin C expression from 30k and 120k cells/scaffold compared to the 10k cells/scaffold (Figure S5c). Interestingly, cell density on the AM 3D scaffolds did not influence the lamin A or C expression (Figure 3b and Figure S5b).

At 5k/cm^2^, cells don’t reach full confluency on 2D films after 7 days, while cells are fully confluent after seeding at 120k/scaffold (Figure 3d). Lamin A and C expression was still much higher on 2D films than on ESP seeded at this high cell density, regardless of the increase in lamin A and C with increased cell density. This shows that even though lamin A and C increase with increased cell density, it does not explain the differences observed here between 2D and 3D cell culture systems.

### 2.5. Less focal adhesions in 3D environments

To understand why lamin A and C expression and the actin organization change in response to dimensionality, we looked at focal adhesion formation of cells cultured on 3D AM- or ESP scaffolds, compared to 2D films. As expected, many large focal adhesions were found on flat films, visualized by paxillin staining, an important focal adhesion protein (**Figure 4a**). Very few and faint focal adhesions were observed in cells cultured in the 3D AM- and ESP scaffolds. Expression of paxillin was also reduced in both 3D AM- and 3D ESP scaffolds, 75% ± 18% (p<0.05) and 57% ± 11% (p<0.05), respectively (Figure 4b).

**Figure 4.**
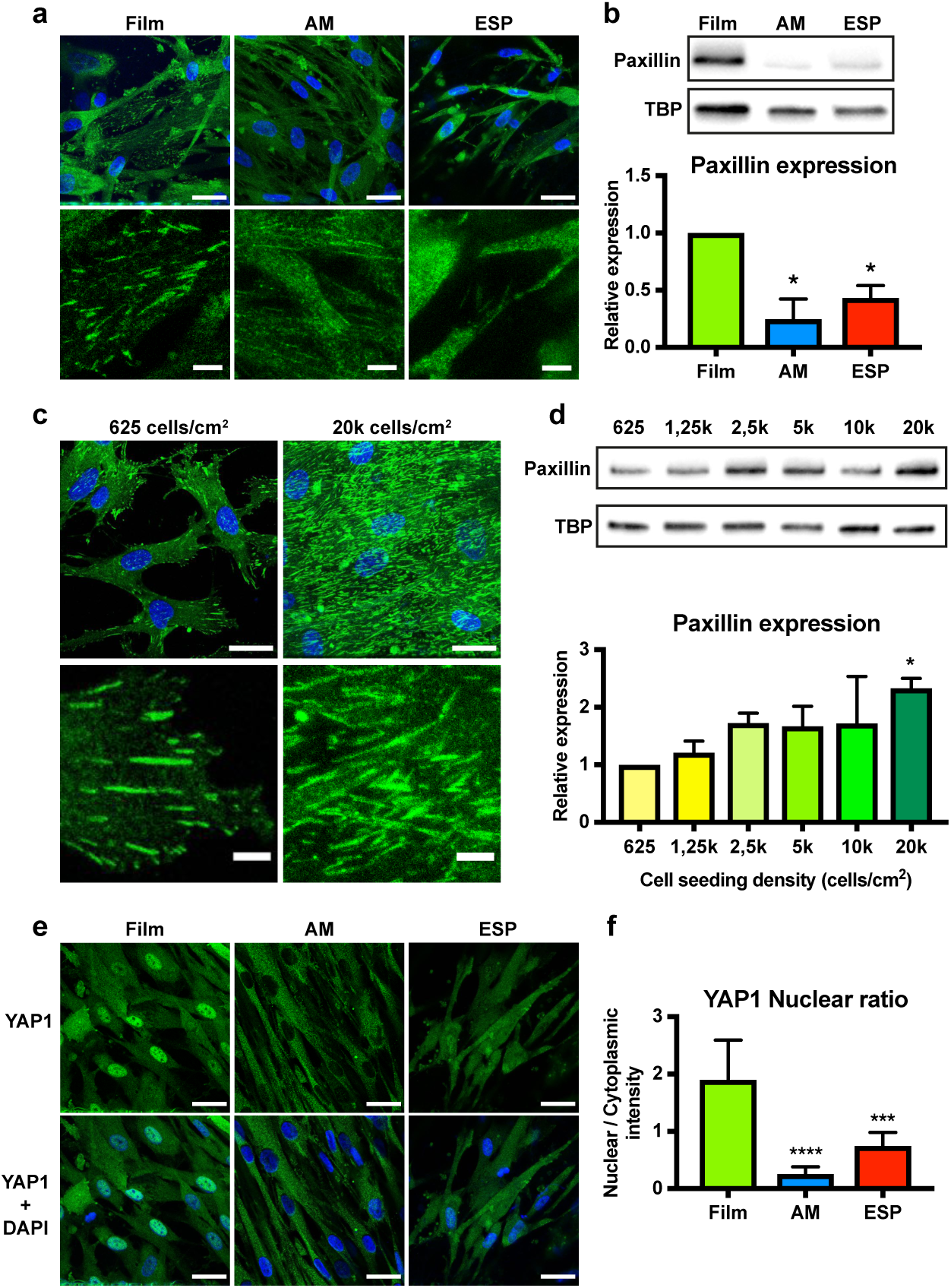
Decreased focal adhesions and cellular tension in 3D. **a,** hMSCs on 2D films, 3D additive manufactured (AM) or electrospun (ESP) scaffolds stained for paxillin (green) and nuclei (blue). The bottom panels are magnifications (5×) of the respective images above. Scale bars represent 30 µm (top panels) and 5 µm (bottom panels). **b,** Paxillin expression in hMSCs cultured on 2D film, 3D AM and ESP scaffolds. TBP is shown as a loading control. Graph shows western blot quantifications of paxillin/TBP, normalized to 2D films, from 4 independent experiments. One-way ANOVA; *p<0.05 compared to 2D films. **c, d,** Paxillin expression in hMSCs seeded on 2D TCP at different cell densities. **c**, Images show paxillin staining (green), with the bottom panels showing magnifications (5×) of the respective images above. Scale bars represent 30 µm (top panels) and 5 µm (bottom panels). **d**, Western blots and quantification of western blots from 3 independent experiments show paxillin expression. TBP is shown as a loading control. Values are normalized to 625 cells/cm^2^. One-way ANOVA; *p<0.05 compared with 1250 cells/cm^2^. **e,** YAP1 staining (green) of hMSCs grown on 2D films, 3D AM and ESP scaffolds. Top panels show YAP1 staining alone, while the bottom panels show YAP1 and nuclei (blue). Scale bars represent 30 µm. **f,** Quantification of the nuclear to cytoplasmic intensity of YAP1 staining in individual cells. Krusal Wallis test; ***p<0.001, ****p<0.0001 compared to films. Total counted cells for films: 27, AM: 28 and ESP: 22 from 4 images.

Less focal adhesions in response to dimensionality correlates to the reduced lamin A and C expression in response to dimensionality. In 2D lamin A and C increased with higher cell seeding densities. To see whether focal adhesion formation also follows the same correlation in 2D, focal adhesion formation and paxillin expression were analyzed in different cell densities in 2D TCP. Indeed, at a lower cell density (625 cells/cm^2^), fewer focal adhesions formed than at higher cell densities (20k/cm^2^) (Figure 4c). Paxillin expression followed this trend, where at 20k/cm^2^ paxillin expression was 2.0 ± 0.43 times higher than at 1.25k/cm^2^ (p<0.05) (Figure 4d). In ESP, where increased cell seeding density also increased lamin A and C, higher cell densities also showed an increase in focal adhesions (Figure S6a). An increase in paxillin expression was visible, but not statistically significantly different (Figure S6b).

Together, these data shows that dimensionality reduces focal adhesion formation, compared to 2D. In addition, a positive correlation between focal adhesions and lamin A and C expression was found, hinting at a possible connection between the two.

### 2.6. Cellular tension reduced in 3D cultures

YAP1 translocates to the nucleus when there is higher cellular tension. ^[33]^ To determine whether cellular tension changes in response to dimensionality, we measured the nuclear translocation of YAP1 in hMSCs cultured on AM or ESP scaffolds, compared to flat films (Figure 4f). From both AM and ESP scaffolds, the nuclear/cytoplasmic ratio of YAP1 was significantly lower (0.3 ± 0.1 (p<0.0001) and 0.7 ± 0.2 (p<0.0001), respectively, compared to that on 2D films 1.9 ± 0.7) (Figure 4e); these low ratios show that more YAP1 remained in the cytoplasm than translocated to the nucleus. The lack of YAP1 nuclear translocation indicates that the dimensionality of the 3D cultures causes a lower cellular tension in hMSCs. This finding is in line with the reduced focal adhesions, actin stress fibers and lamin A and C expression in response to dimensionality.

Previous work proposed that YAP1 regulates lamin A and C. ^[6, 34^] To test if YAP1 plays a role in lamin A and C expression, we knocked down YAP1 in hMSCs cultured on TCP and determined the levels of lamin A and C expression. No difference in lamin A or C expression was found between scrambled control and YAP1 knock-downs. This shows that lamin A and C is not regulated through YAP1 (Figure S7) and indicates that the lower lamin A and C levels in the 3D scaffolds are not due to a lack of YAP1 activity in the 3D scaffolds.

### 2.7. Zyxin influences lamin A and C expression and actin organization

The dimensionality caused low lamin A and C levels and few focal adhesions and actin stress fibers. We next determined whether the focal adhesions are involved in controlling the low lamin A and C expression and resulting changed actin network. Zyxin is known to be critical for focal adhesion formation and the connection to the actin cytoskeleton ^[35, 36]^, and was therefore chosen as target to diminish the focal adhesion formation. We knocked down zyxin in hMSCs using two different ZYX-shRNAs (Figure S8), cultured the cells on TCP, then measured lamin A and C expression and evaluated the actin network.

Lamin A was reduced 92 ± 5% (p<0.01) and 71 ± 13% (p<0.05) and lamin C reduced 86 ± 11% and 76 ± 19% by the two ZYX-shRNAs, respectively, compared to the scrambled control (**Figure 5a**, Figure S9). In the ZYX knockdowns, less actin was observed in the cell center and over the nucleus, while more F-actin was found in the cell periphery compared to that of the scrambled control (Figure 5c and d), a localization pattern similar to the LMNA knockdown and cells cultured on ESP scaffolds (Figure 1). The reduction of stress fibers after ZYX knockdown is in line with previous reports. ^[35, 36]^

**Figure 5.**
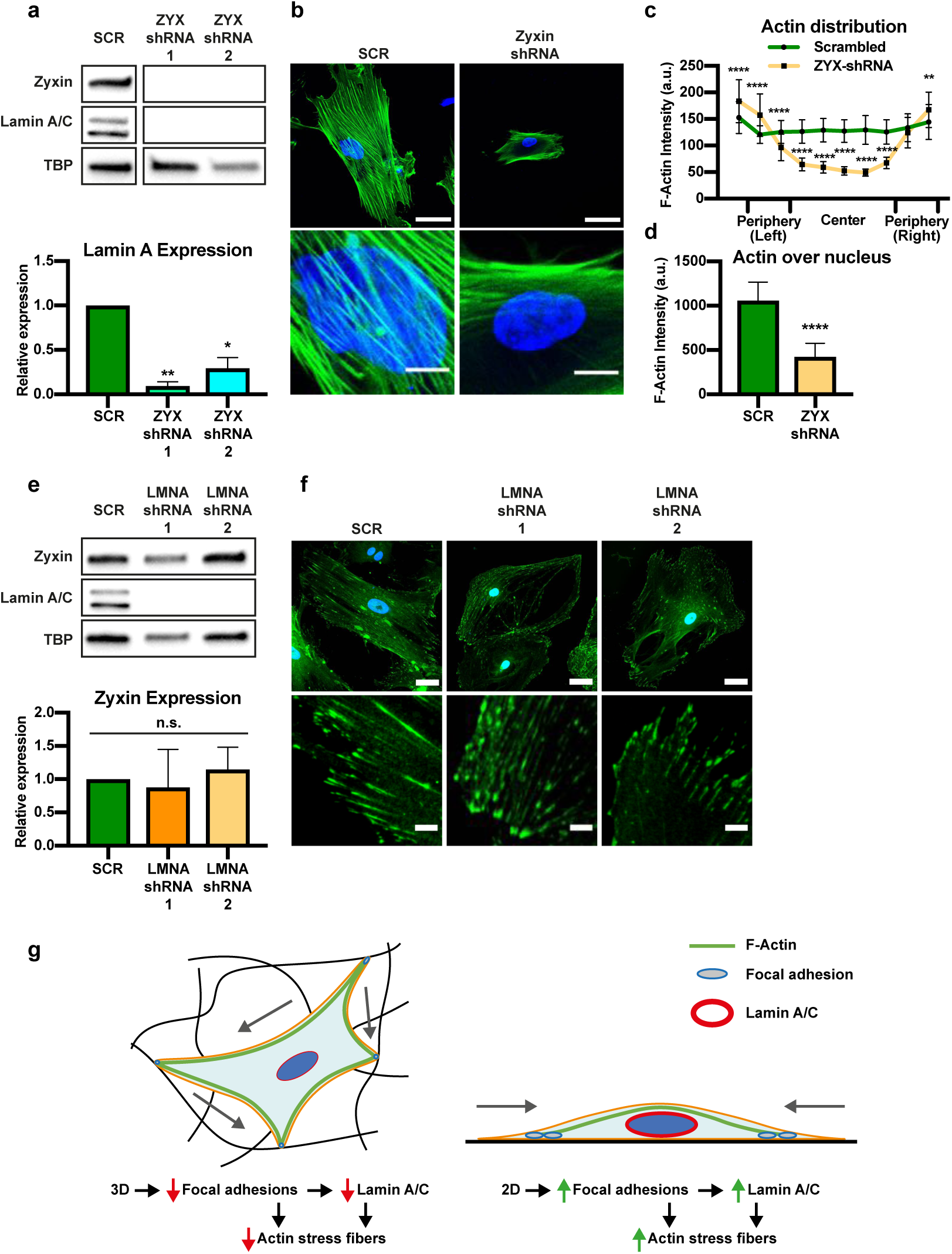
Zyxin influences lamin A and C expression, but not vice versa. **a, e,** Western blots of zyxin and lamin A and C expression in hMSCs transduced with scrambled or two different ZYX-shRNAs (a) or two different LMNA-shRNAs (e). Graphs show the quantification of lamin A/TBP (a) and zyxin/TBP (e) expression averaged from 4 biological replicates, and normalized to expression from SCR-shRNA. Error bars represent mean ± SD. One-way ANOVA; **p<0.01 compared to SCR. **b,** hMSCs transduced with scrambled or ZYX-shRNA, stained for F-actin (green) and nuclei (blue). The bottom panels show 5×magnifications of the respective images above. Scale bars represent 50 µm (top) and 10 µm (bottom). **c,** Quantification of the F-actin intensity distribution in hMSCs transduced with scrambled or ZYX-shRNA. n=33 cells analyzed for SCR and n=32 for ZYX-shRNA from 3 biological replicates. Two-way ANOVA; ****p<0.0001. **d,** Quantification of F-actin intensity that overlaps with nuclei staining in hMSCs transduced with scrambled or ZYX-shRNA cultured on 2D TCP. n=25 cells analyzed for SCR and n=26 for ZYX from 3 biological replicates. Mann-Whitney test, ****p<0.0001. **c, d,** Error bars represent mean ± 95% CI. **f,** hMSCs transduced with scrambled or LMNA-shRNAs stained for zyxin (green) and nuclei (blue). Bottom panels show 5× magnifications of the respective images above. Scale bars represent 50 µm (top panels) and 10 µm (bottom panels). **g,** Model showing differences in actin fiber formation, focal adhesion and lamin A and C in 3D (left) v 2D (right.) In 3D, cellular tension forces (grey arrows) are distributed in multiple directions and focal adhesions (light grey ovals) stay small and few. The reduction of focal adhesions then leads to a reduction in lamin A and C expression (red oval). Together, small and few focal adhesions and low lamin A and C expression then lead to the lack of actin stress fibers (green lines) through the cell center. In 2D, cellular tension forces (grey arrows) are distributed along a single plane, allowing for the build-up tension and large focal adhesions (light grey ovals). This leads to an increase in lamin A and C expression (red oval). Large focal adhesions and high lamin A and C expression then enable the formation of actin stress fibers (green lines) through the cell center.

Zyxin also greatly reduced the number and intensity of paxillin positive focal adhesions, although it could still be visualized (Figure S10), also in line with previous reports. ^[37]^ Paxillin is an important protein for the downstream signaling of focal adhesions. ^[38]^ To test if lamin A and C expression are influenced by paxillin, we knocked paxillin down. No difference was observed in lamin A or C expression, demonstrating that the reduction of lamin A and C following zyxin knock down does not occur through paxillin (Figure S7).

To test whether the influence of zyxin on lamin A and C expression is bidirectional, we looked at zyxin expression in LMNA knock-downs. Zyxin expression and focal adhesion formation were similar in LMNA knockdowns and scrambled controls (Figure 5e and f), demonstrating that lamin A and C expression does not affect zyxin levels. Furthermore, these data suggest that the faint focal adhesions we observed from hMSCs on 3D scaffolds (Figure 4a) were not due to the low lamin A and C expression.

Interestingly, in both ZYX and LMNA knockdowns, YAP1 was still located mainly in the nucleus (Figure S11). This indicates that the lack of YAP1 nuclear localization in 3D is through a mechanism independent of lamin A and C or zyxin.

### 2.8. A model for actin fiber formation mediated by lamin A and C and zyxin in 2D v. 3D

Our data comes together in the model depicted in Figure 5g. In 3D, dimensionality prevents focal adhesion formation, which leads to a decrease in lamin A and C. The lack of both zyxin and lamin A and C then shapes the actin network and does not allow for the formation of actin stress fibers through the cell center. In 2D, the lack of dimensionality allows for the formation of many large focal adhesions, leading to an increase in lamin A and C. High zyxin and lamin A and C expression then allow for the formation of high cellular tension and large actin stress fibers. This mechanism is independent of both YAP1 and paxillin.

## 3. Discussion

In this study, we show that hMSCs cultured in 3D ESP or AM scaffolds exhibit low cellular tension, even on stiff materials. This was visualized by a decrease in focal adhesions, lamin A and C expression, YAP nuclear localization and actin stress fibers on 3D ESP and AM scaffolds, compared to 2D films of the same material. We show that zyxin influences lamin A and C expression, but not vise-versa. Both zyxin and lamin A and C play a role in the formation of stress fibers in the cell center, and are partly responsible for the different actin organization in 3D.

Fraley et al. also showed a decrease in focal adhesions in response to dimensionality in or on hydrogels. ^[26]^ Also, in a 3D *in vivo* environment, focal adhesions were found to be very different in shape and composition from 2D cultures. ^[39]^ How dimensionality initiates a difference in the focal adhesion protein zyxin and focal adhesion formation has not been studied here and remains unclear. Potential mechanisms include a difference in force distribution. On concave surfaces, focal adhesion formation and lamin A and C have been shown to decrease when compared to flat surfaces, while they increase on convex surfaces.^[40]^ This indicates that the force distribution within the cell and the angle at which it is connected to the environment (partly) determines focal adhesion formation and lamin A and C expression. We hypothesize that because of the dimensionality, forces are not distributed over the nucleus like in 2D, but more throughout the whole cell and along the cell periphery, which could explain the more peripheral actin network in 3D environments. The knock down of lamin A and C could mimic what happens in a 3D environment, where forces can no longer be distributed to the nucleus, which leads to the lack of actin stress fibers near the nucleus. Indeed, stress fibers and focal adhesions can still be found in the LMNA knock downs, but are then located almost exclusively in the cell periphery (Figure 2b, 2c, 5f).

We found that YAP1 was excluded from the nucleus on ESP and AM scaffolds, in contrast to 2D films. However, in LMNA and ZYX knockdowns, YAP1 was still located in the nucleus. The opening of nuclear pores due to force on the nucleus has been shown to allow YAP1 to enter the nucleus. ^[33]^ A possible explanation is that because of the lower nuclear stiffness in both LMNA and ZYX knockdowns (due to lack of lamin A and C), YAP1 can enter the nucleus even if the cell is under less tension. ^[33]^ In 3D, the nucleus might be under less tension, as forces are distributed in all 3 dimensions, whereas in 2D forces are mostly concentrated in two directions, pushing down on the nucleus. ^[33]^ This lack of tension on the nucleus could potentially result in avoiding YAP1 to enter the nucleus in 3D.

Here, we have presented a novel link between lamin A and C and the actin cytoskeleton, and lamin A and C and focal adhesions, specifically zyxin. We have found that this process is not mediated by YAP1. Other mechanosensitive (co-)transcription factors could be inhibited to look at the effect on lamin A and C and zyxin, to find the signaling pathway involved in orchestrating the differences in expression in 3D environments.

Decoupling individual variables in 2D vs 3D in stiff materials remains challenging. Local and bulk material properties can change when creating scaffolds, potentially affecting cell behavior. Also, ECM produced by cells can be influenced, directly changing the environment. Using hydrogels, 2D vs 3D can be more strictly controlled. We show here for the first time that in stiff materials, the same reduction in cellular tension is observed when cells are cultured in a 3D environment. ^[26]^ This can have important implications, as it is known that differentiation is highly influenced by cellular tension, YAP1, lamin A and C, focal adhesions and actin stress fibers. ^[22–25]^ Our data shows that using stiff materials might not be sufficient to induce high cellular tension when cells are cultured in 3D. Also, soft tissue constructs could still be made of relatively stiff materials, if the dimensionality decreases cellular tension to still allow for effective differentiation to soft tissue lineages. An optimized microenvironment could include different concave or convex surfaces, different pore geometries or mechanical loading to optimize the cellular tension and differentiation of hMSCs in 3D tissue engineering constructs.

## 4. Conclusion

Here, we study the effect of dimensionality on cellular tension of hMSCs in two common 3D tissue engineering constructs: ESP and AM scaffolds. We demonstrate that dimensionality causes fewer stress fibers, fewer focal adhesion, lower lamin A and C expression, less YAP1 nuclear localization. Cell density influences lamin A and C expression and focal adhesion formation in 2D and 3D, but it is not responsible for the observed differences between 2D and 3D, further proving that dimensionality is an important environmental factor. Lamin A and C or zyxin knock down result resulted in fewer stress fibers in the cell center and over the nucleus. Zyxin knock down also reduced lamin A and C expression, but not vice versa, showing that this signal chain starts from the outside of the cell. Taken together, our study shows dimensionality changes the actin cytoskeleton through lamin A and C and zyxin and decreases cellular tension, even on stiff materials. This can have important implications for future scaffold design, as proteins involved in cellular tension also play key functions in hMSC differentiation.

## 5. Experimental Section

### Scaffold production

Poly(ethylene oxide terephthalate) and poly(butylene terephthalate) random block co-polymer (PEOT/PBT, 300PEOT55PBT45, PolyActive™) with 300 Da PEO and a PEOT/PBT weight ratio of 55/45 was acquired from PolyVation. Polystyrene (PS, 350 kDa) was acquired from Sigma-Aldrich. PEOT/PBT or PS films were produced by melting the polymer in a circular 23-mm mold between two silicon wafers (Si-mat, Kaufering, Germany) and two hot plates under slight pressure (∼100 kg) to ensure films were fully flat. PEOT/PBT was processed at 180 °C and PS at 210 °C.

AM scaffolds were produced by means of screw-extrusion–based fused deposition modeling (FDM) (Bioscaffolder SYSENG, Germany). The FDM extrusion is controlled by the screw rotation and assisted by N2 (5 bar) gas pressure allowing fine control over deposition of the molten polymer. The manufacturing of the 20×20×4 mm scaffolds was achieved following a layer-by-layer manufacturing with 90° rotation between deposited layers. The 3D scaffold CAD models were uploaded into PrimCAM software (Primus Data, Switzerland) and the deposition patterns were calculated. The fiber spacing, defined as the distance between successive fibers in the same layer was defined as 650 µm, the layer thickness was set to 170 µm, and the fiber diameter obtained was according to the nozzle diameter used, the polymer selected and the processing parameters. The parameters that influence the production of the 3D scaffolds are: temperature, screw rotation, deposition velocity. PEOT/PBT or PS pellets were loaded in the reservoir and molten at a temperature of 195 °C or 220 °C, respectively. The screw rotation for the polymers was 200 rpm. The molten polymer was extruded through a nozzle with G25 (I.D. = 250 µm). The deposition velocity was optimized to 20 and 200 mm·min^-1^ for PS and PEOT/PBT, respectively.

To produce ESP scaffolds, 20% (w v^-1^) 300PEOT55PBT45 was dissolved in a mixture of 70% chloroform (Sigma-Aldrich) and 30% 1.1,1.3,3.3-Hexafluoro-2-propanol AR (HFIP; Bio-Solve) overnight under agitation at room temperature. Electrospinning was done on a mandrel (diameter: 19 cm) rotating at 100 rpm to produce many scaffolds at the same time under exactly the same conditions. The following conditions were maintained: 15 cm working distance, 1 ml h^-1^ flow rate, 23–25 °C and 40% humidity. The mandrel was charged between - 2 and -5 kV and the needle between 10–15 kV. ESP scaffolds were spun on aluminum foil over a polyester mesh with 12-mm holes. After spinning, ESP scaffolds with a diameter of 15 mm were punched out and scaffolds were removed from the aluminum foil. This method yielded ESP scaffolds of 12 mm with a 1.5-mm supporting ring of polyester mesh around them to improve handleability. Fibers were deposited randomly with a diameter of 0.99 ± 0.18 µm, creating mats of approximately 50-µm thick.

Before cell culture, films, AM and ESP scaffolds were sterilized with 70% ethanol for 15 min and dried until visually dry. The films and scaffolds were then coated with 1 mg ml^-1^ rat-tail collagen I solution for 16 h at 37 °C. After coating, they were washed twice with water and air-dried before cell seeding.

### Cell culture

For expansion, hMSCs were cultured on TCP at 1000 cells/cm^2^ in αMEM + 10% fetal bovine serum (basic medium) (Thermo-Fisher Scientific), until 70–80% confluent. All experiments were performed at passage 5. Films were punched with a diameter of 22 mm and cultured in non-treated, 12-well plates at 5k cells/cm^2^, unless stated otherwise. ESP scaffolds were 15 mm and cultured in 24-well plates at 30k cells/scaffold, unless stated otherwise, with a rubber O-ring (outer diameter 15 mm, inner diameter 12 mm) to prevent the scaffolds from floating and to cover the polyester ring on which the scaffolds were produced. AM scaffolds were square blocks of 5×5×3 mm (width, length, height) and cells were seeded in a drop of 50 µl containing 400k cells, unless stated otherwise. Two hours after seeding, the AM scaffold was flipped upside down to increase cell distribution. After a total of 4 h after seeding, scaffolds were transferred to a non-treated, 12-well plate for further culture. Film and scaffold cultures were done in basic medium supplemented with 1 ng ml^-1^ FGF-2 (Neuromics), 200 µM L-Ascorbic acid 2-phosphate (Sigma-Aldrich) and 100 U ml^-1^ penicillin-streptomycin, and were harvested after 7 d of culture, unless stated otherwise.

### Protein isolation and western blot

Proteins were isolated from cells cultured on films or scaffolds in RIPA buffer (Sigma-Aldrich), supplemented with cOmplete™, Mini, EDTA-free Protease Inhibitor Cocktail (Sigma-Aldrich). To get sufficient proteins, 6–12 films, 15–20 ESP scaffolds or 2–4 AM scaffolds were mixed into 300–400 µl lysis buffer for a single protein isolate. Experiments were repeated 3 or 4 times for replicates. For TCP and films, surfaces were scraped with cell scrapers to ensure cell lysis. AM scaffolds were cut into four smaller pieces and submerged in lysis buffer. ESP scaffolds were removed from the polyester supporting ring and submerged in lysis buffer. Samples were spun down at 10.000*g*, and the supernatant was used for further processing.

Protein quantification was done using the Pierce BCA protein assay kit (Thermo Fisher Scientific). Twenty micrograms of protein was incubated with laemmli loading buffer (Bio-Rad) and 10% 2-Mercaptoethanol (Sigma-Aldrich) for 5 min at 95 °C and loaded into a 4– 15% polyacrylamide gel (Bio-Rad). Proteins were transferred to a 0.45 µm PVDF membrane (Bio-Rad) using the semi-dry transfer method. Membranes were blocked for 1 h with 5% fat free milk powder (Bio-Rad) in TBS + 0.05% tween-20 (Sigma-Aldrich). All primary antibody incubations were performed overnight at 4 °C in blocking buffer. All antibodies (lamin A and C: ab108595; paxillin: ab32084; zyxin: ab58210; YAP1: ab52771; TBP: ab51841) were ordered from Abcam and diluted 1/1000, except for YAP1 which was diluted 1/500. Blots were subsequently incubated with 0.33 µg ml^-1^ goat-anti-rabbit or mouse horseradish peroxidase (Bio-Rad) in blocking buffer for 1 h at room temperature. Protein bands were then visualized using Clarity Western ECL (Bio-Rad). Quantifications of band intensity were done in Fiji using the gel quantification tool.

### Immunofluorescence and imaging

Prior to staining, hMSCs were fixed in 3.6% (v v^-1^) paraformaldehyde (Sigma-Aldrich) in PBS for 20 min at room temperature. Cells were permeabilized and blocked in 2% bovine serum albumin (BSA) (VWR) and 0.1% triton X (VWR) in PBS for 1 h at room temperature. Antibodies mentioned above, in the same dilution, were incubated overnight at 4 °C in 2% BSA and 0.05% tween-20 in PBS (incubation buffer). Secondary antibodies goat-anti-rabbit or mouse Alexa Fluor 488 (Thermo Fisher Scientific) were then incubated overnight at 4 °C in incubation buffer. Instead of antibody staining, F-actin staining was done with phalloidin Alexa Fluor 488 (Thermo Fisher Scientific) at room temperature for 20 min in PBS+0.05% tween-20. Nuclei were stained with DAPI (Sigma-Aldrich). Images were taken on a widefield fluorescence microscope, or confocal microscope. To allow for quantitative comparison between images, all images within an experiment were taken with the same settings and on the same day.

Actin quantification was done using a custom-built Fiji macro *(paper in submission)*. A line was drawn through the cell perpendicular to the long axis of the cell, and the intensity over that line was measured. The intensity distribution was divided into 10 equal bins to correct for differences in cell size. The total F-actin intensity overlapping with the nucleus was also measured.

For YAP1 quantification, the nucleus and an area in the cytoplasm were selected and the overall pixel intensity in each area was measured using Fiji. The ratio between nuclear and cytoplasmic intensity was then calculated per cell.

### Migration assay

As a functional read-out for nuclear stiffness ^[29]^, hMSCs were allowed to migrate directly from films or ESP scaffolds through a Fluoroblok 24-well transwell with 3-µm or 8-µm pores. hMSCs were first cultured on 6-mm films or ESP scaffolds for 7 days in standard scaffold culture conditions (see ‘Cell culture’) and then transferred upside down on the transwell membrane. αMEM without serum was added to the top compartment and basic medium was added to the bottom compartment, to induce cell migration. A screw and nut were placed on top of the film or ESP scaffold as a weight to ensure proper contact between the substrate and transwell. After fixation (see ‘Immunofluorescence and imaging’), the film or ESP scaffold was removed, and the transwells were stained with Syto14 (Thermo Fisher Scientific). The whole transwell was imaged, top and bottom, and cells were quantified using particle analysis in Fiji.

### Lentiviral production for shRNA delivery

To deliver shRNA for gene knock-down, we produced lentiviruses using TRC pLKO.1 constructs from Dharmacon. We used the following clone IDs for LMNA: TRCN0000061833 and TRCN0000061836; ZYX: TRCN0000074204 and TRCN0000074205; PXN: TRCN0000123134 and TRCN0000123136; YAP1: TRCN0000107265 and TRCN0000107266. To produce the lentiviral particles, human embryonic kidney 293FT (HEK) cells were seeded at 60k cells/cm^2^ in a TCP dish in DMEM + 10% FBS. The next day, HEK cells were transfected with pMDLg pRRE, pMD2.G, pRSV Rev (Addgene) and one of the pLKO.1 plasmids containing the shRNA, using lipofectamine 2000 (Thermo Fisher Scientific) in ratio of 5:1 (µl:µg of DNA). After overnight incubation, the medium was changed to basic medium for hMSCs. Viral particles were harvested after 24 and 48 h, and filtered through a 0.45-µm filter.

The day before transduction, hMSCs were thawed at 1k cells/cm^2^ in a 10-cm dish. 3 milliliters of unconcentrated virus was added to the dish and incubated overnight. The medium was changed for basic medium the following day, and the medium was changed for basic medium + 2 µg ml^-1^ puromycin at 48–72 h later. Cells were treated for 72 h with puromycin. At 9–10 days after initial thawing, cells were passaged at 1k cells/cm^2^ and cultured for 7 days in basic medium before protein harvest or fixation.

### Statistical analysis

The number of biological replicates and repeated experiments are indicated in the figure captions, as well as the statistical test used. All individual films and scaffolds used in one experiment were randomly assigned to an experimental group. Cells that were imaged for quantification were also randomly picked. All data was tested for normal distribution using the Shapiro-Wilk test. To test the significance of relative expression data with one comparison, a two-tailed ratio t-test was performed, or the Mann-Whitney test as non-parametric equivalent. For relative expression data with multiple comparisons, the log of each value was used for a repeated measures ANOVA, with Tukey’s post hoc to compare individual groups. For non-relative comparisons, a One-way ANOVA with Tukey’s post hoc, or Two-way ANOVA with Sidak’s post-hoc was used, or the Kruskal-Wallis test with Dunn’s post hoc as non-parametric equivalent. Statistical significance was set at p<0.05.

## Acknowledgements

We would like to thank Jos Broers for valuable discussions on the lamin A and C data. We are grateful to the European Research Council starting grant “Cell Hybridge” for financial support under the Horizon2020 framework program (Grant #637308). Some of the materials used in this work were provided by the Texas A&M Health Science Center College of Medicine Institute for Regenerative Medicine at Scott & White through a grant from NCRR of the NIH (Grant #P40RR017447).

## Data Availability

Data is avaialable upon request

---

**Figure S1.**
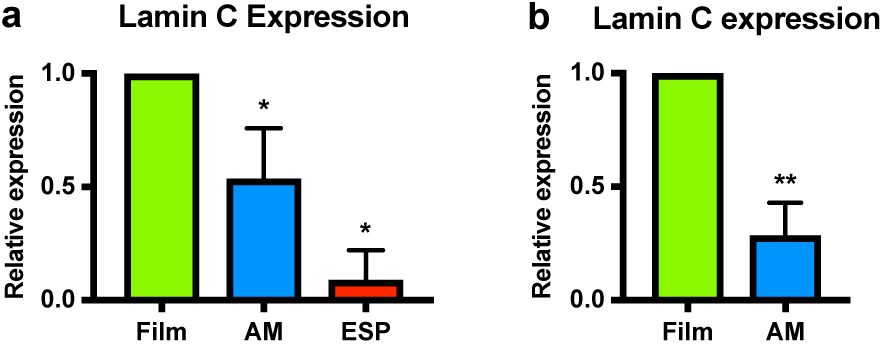
Reduced lamin C expression in 3D. **a,** Lamin C expression in hMSCs grown on PolyActive films, additive manufactured- or electrospun scaffolds. Graph depicts average expression of lamin C/TBP normalized to films, quantified by western blots from 4 independent experiments. One-way ANOVA; * p<0,05, compared with films. **b,** Lamin A and C expression of hMSCs cultured on PolyStyrene films or additive manufactured scaffolds. Graph shows the average expression of lamin A/TBP normalized to films, quantified by western blot from 3 independent experiments. Ratio paired t-test; ** p<0,01. Error bars represent mean±95% CI.

**Figure S2.**
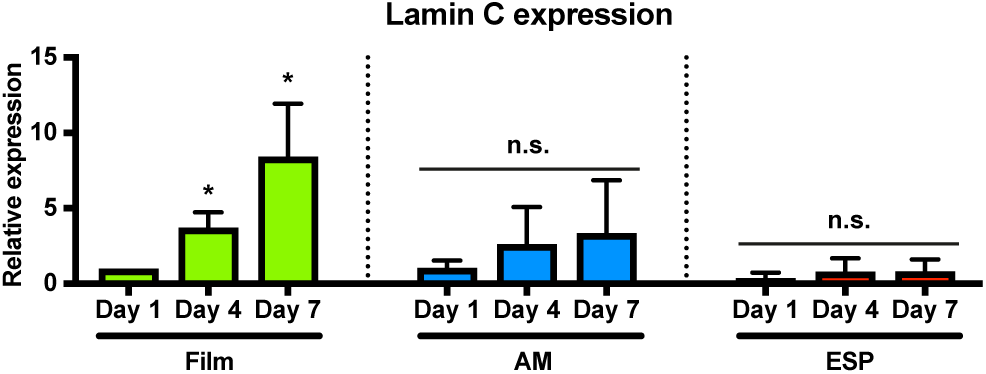
Lamin C expression increases over time in 2D, but not in 3D. **a,** Lamin C expression on day 1, 4 and 7 after seeding hMSCs on films, additive manufactured- or electrospun scaffolds. Graph depicts average expression of lamin C/TBP normalized to films day 1, quantified by western blots from 3 independent experiments. One-way ANOVA; * p<0,05 compared with film day 1. Error bars represent mean±SD.

**Figure S3.**
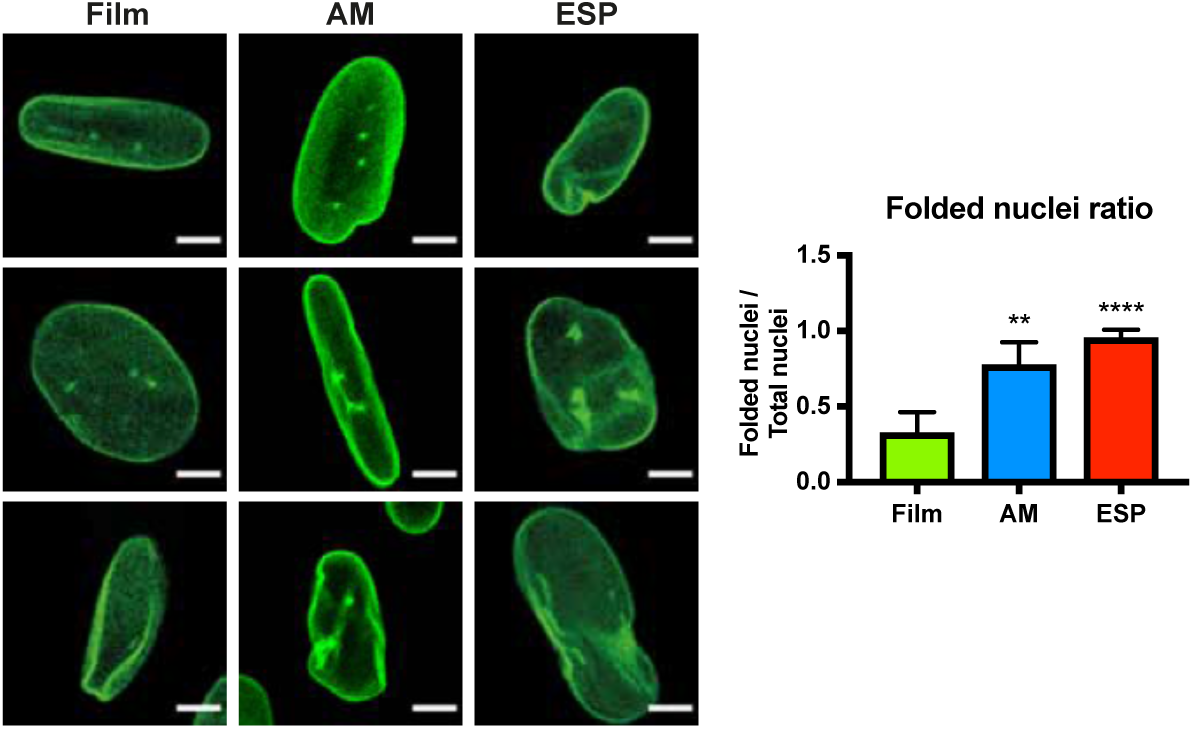
More flexible nuclei in 3D. Representative images of examples of folded and non-folded nuclei of hMSCs grown on films, additive manufactured scaffolds or electrospun scaffolds stained with Lamin A and C (green), scalebars 5µm. The graph shows the corresponding quantification. Total counted nuclei for films: 491, AM: 159, ESP: 79, in 8, 11 and 10 different images, respectively, from 2 independent experiments. Error bars represent mean±95% CI. Kruskal Wallis test, ** p<0,01, **** p<0,0001 compared to films.

**Figure S4.**
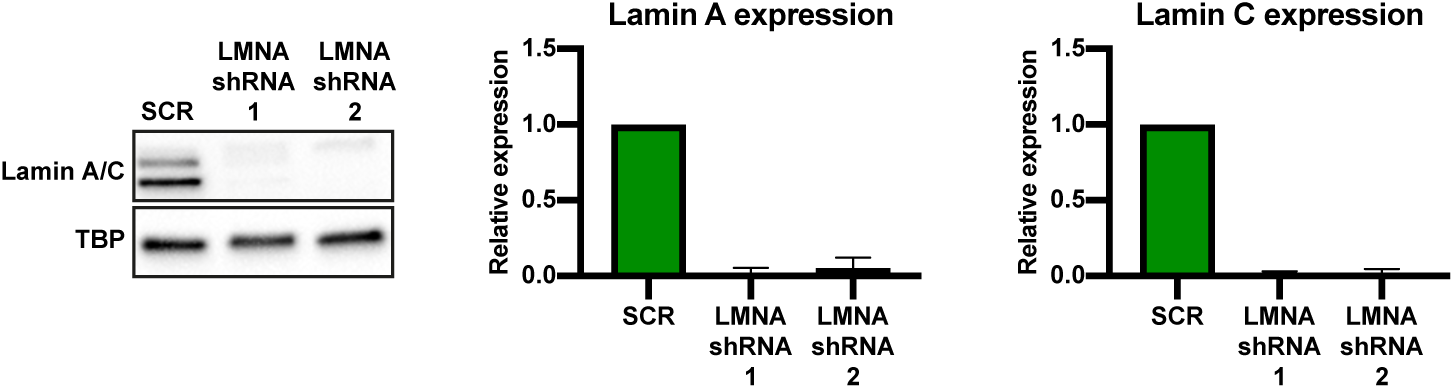
Proof of LMNA knock downs. Quantification by western blot of Lamin A and C of hMSCs transduced with scrambled-, or 2 different LMNA shRNA’s. TBP is shown as loading control. Graphs show the average expression of lamin A or C/TBP, normalized to SCR of 4 biological replicates. Error bars represent mean±SD.

**Figure S5.**
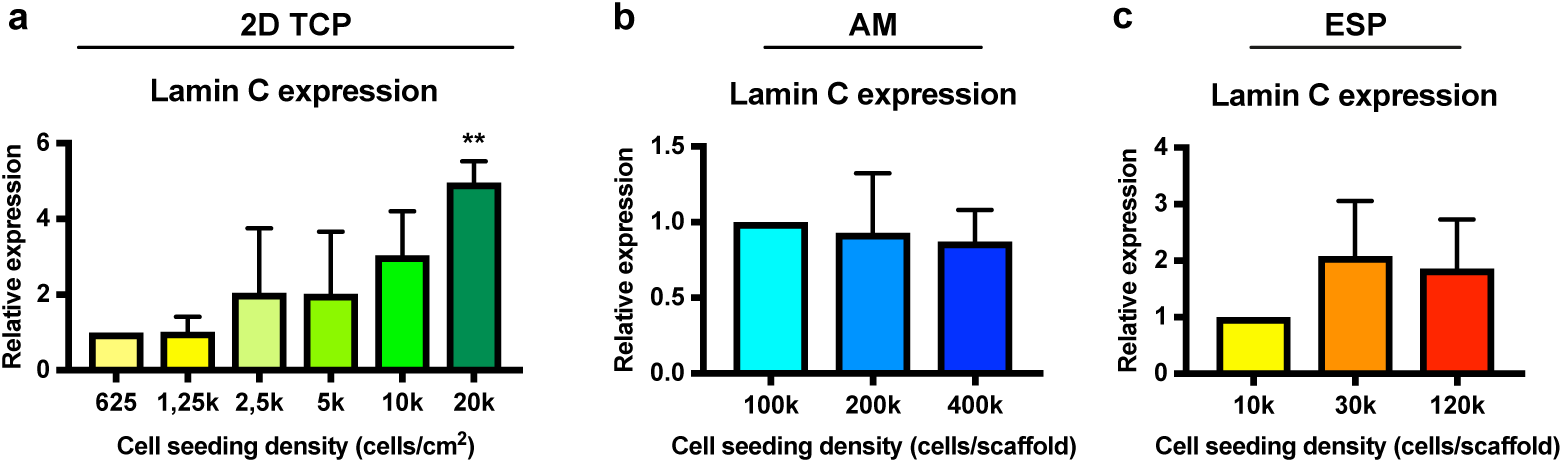
Increased expression of lamin C with increased cell seeding density in 2D. **a,** Lamin C expression of hMSCs seeded on TCP in varying densities. Values are nomalized to 625 cells/cm^2^. One-way ANOVA; * p<0,05 compared with 1250cells/cm2. **b, c,** Lamin C expression of hMSCs on additive manufactured scaffolds (b), or ESP (c), seeded in three different cell densities. Values normalized to 100k (b) or 10k (c) cells/scaffold. **a, b, c,** Graph shows average expression of lamin C/TBP, quantified by western blots from 3 independent experiments. Error bars represent mean±SD

**Figure S6.**
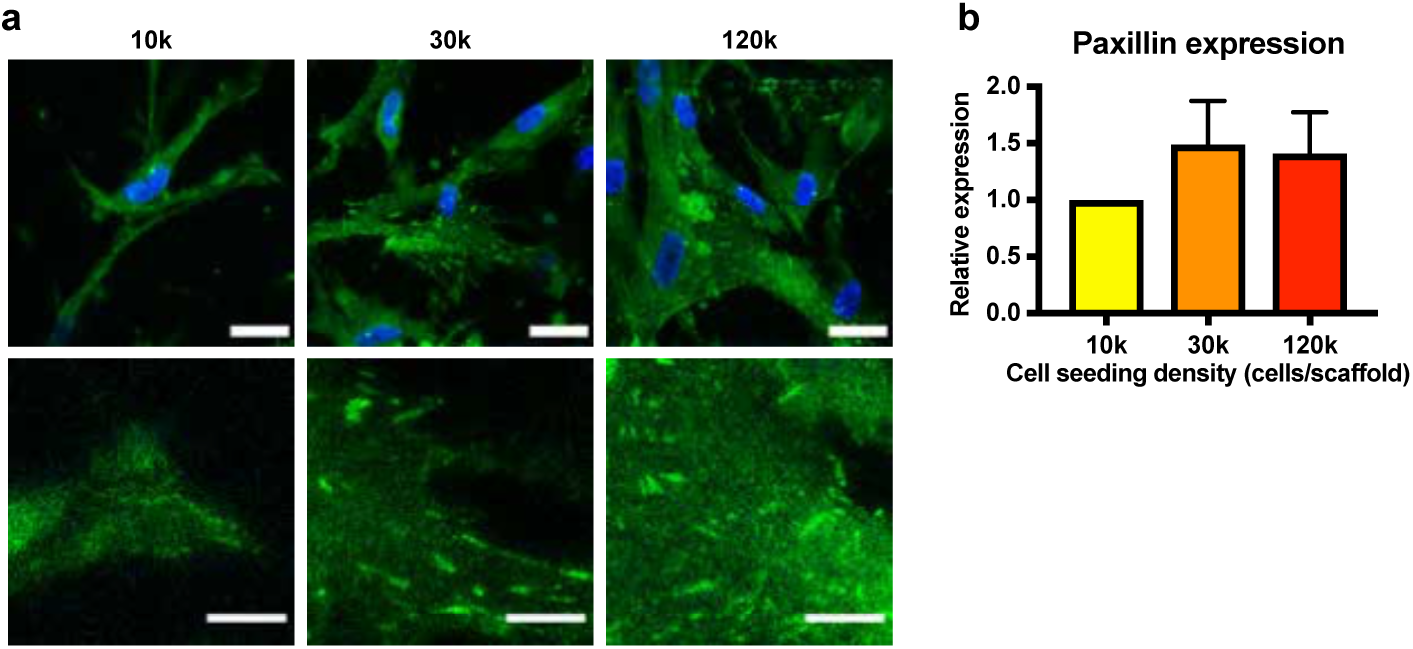
Increased focal adhesion formation with higher cell seeding density on ESP. **a,** Paxillin (green) and nuclei (blue) staining of hMSCs seeded on electrospun scaffolds in 3 different cell densities. Bottom panel shows a 4x magnification of the respective image above. Scalebar 30µm (top panel) and 10µm (bottom panel). **b,** Quantification paxillin expression by western blot of four independent experiments, of hMSCs seeded in different cell densities on electrospun scaffolds. Graph shows Paxillin/TBP, normalized to 10k. Error bars represent mean±SD.

**Figure S7.**
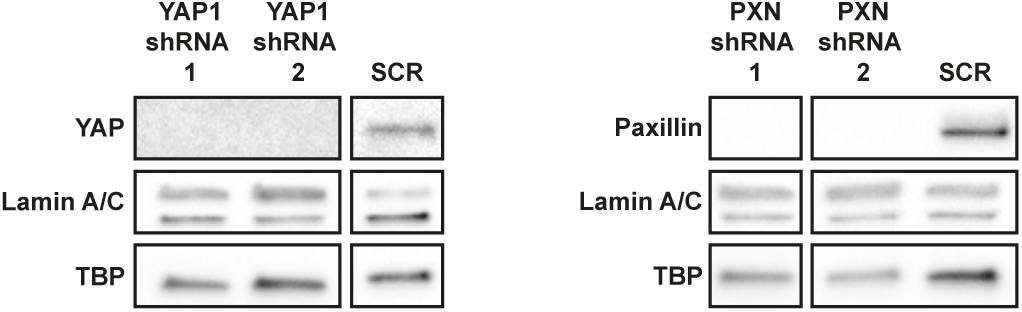
Lamin A and C expression is not regulated by YAP1 or Paxillin. Western blot of hMSCs knocked down for YAP1 (left), or paxillin (right). TBP is shown as loading control. Representative western blot of two independent experiments.

**Figure S8.**
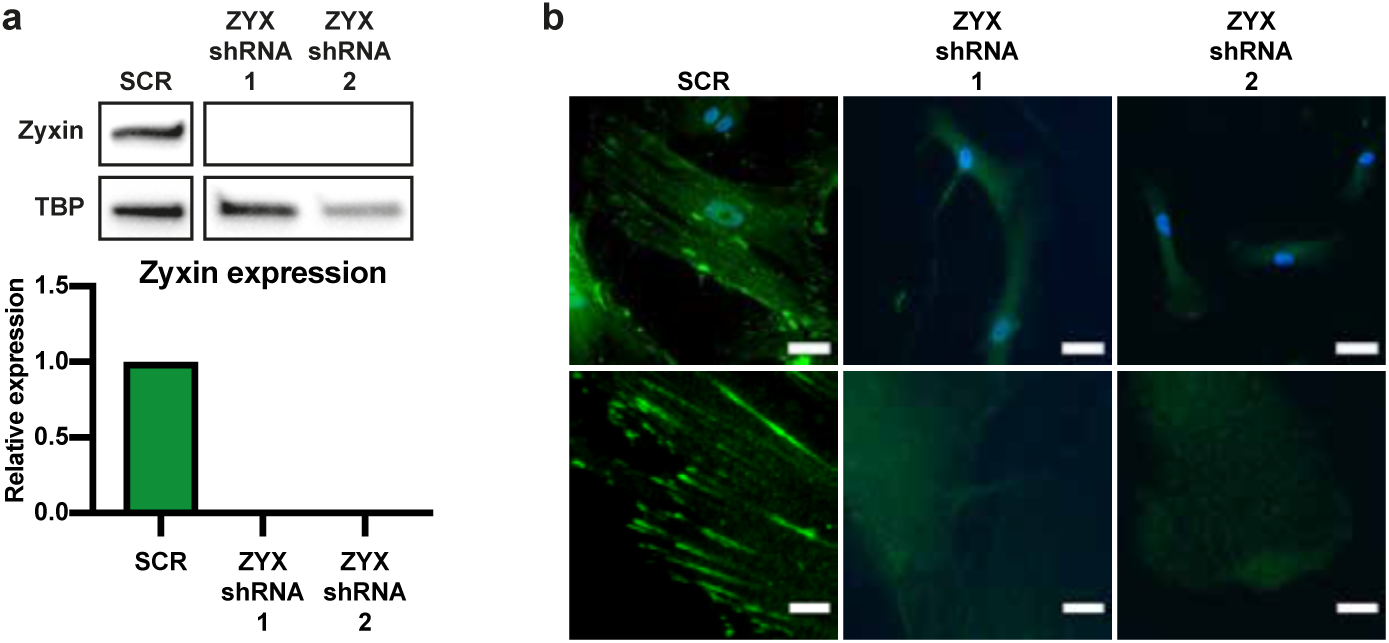
Proof of zyxin knockdowns. **a,** Quantification by western blot of zyxin of hMSCs transduced with scrambled-, or 2 different ZYX shRNA’s. TBP is shown as loading control. Graphs show the average expression of Zyxin/TBP, normalized to SCR of 4 biological replicates. **b,** Staining of zyxin (green) and nuclei(blue) in scramble- or zyxin knockdown-hMSCs. Bottom panel shows a 5x magnification of the respective image above. Scalebars 50µm (top panel) and 10µm (bottom panel).

**Figure S9.**
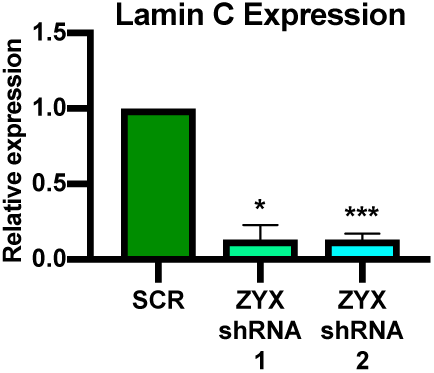
Decreased lamin C expression in cells knocked down for Zyxin. Western blot of hMSCs transduced with scrambled- or two different ZYX-shRNA’s. Graph shows the lamin C/TBP expression normalized to SCR, averaged from 4 biological replicates. Error bars represent mean±SD. One-way ANOVA; * p<0,05, *** p<0,001, compared to SCR.

**Figure S10.**
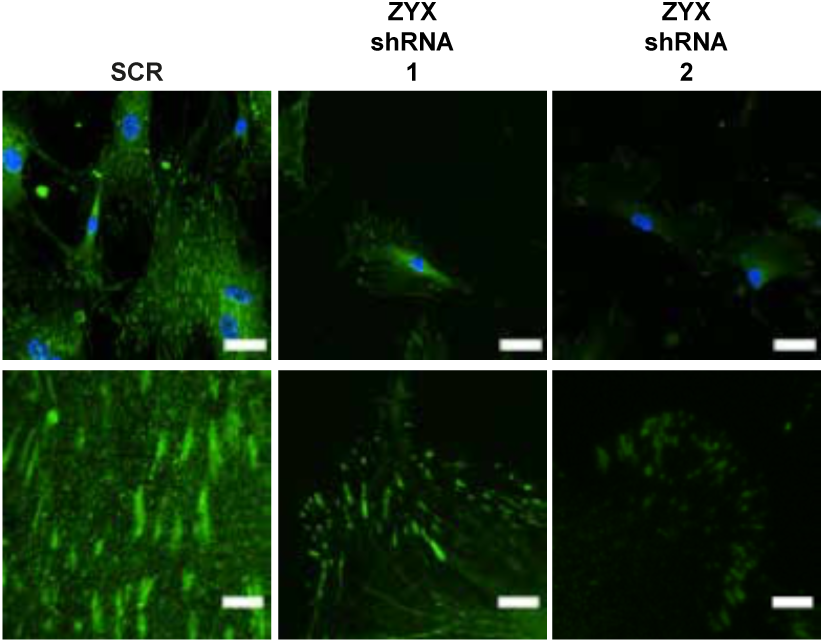
Reduction of paxillin positive focal adhesions in ZYX knockdowns. Staining of paxillin (green) and nuclei (blue) in scramble- or zyxin knockdown-hMSCs. Bottom panel shows a 5x magnification of the respective image above. Scalebars 50µm (top panel) and 10µm (bottom panel).

**Figure S11.**
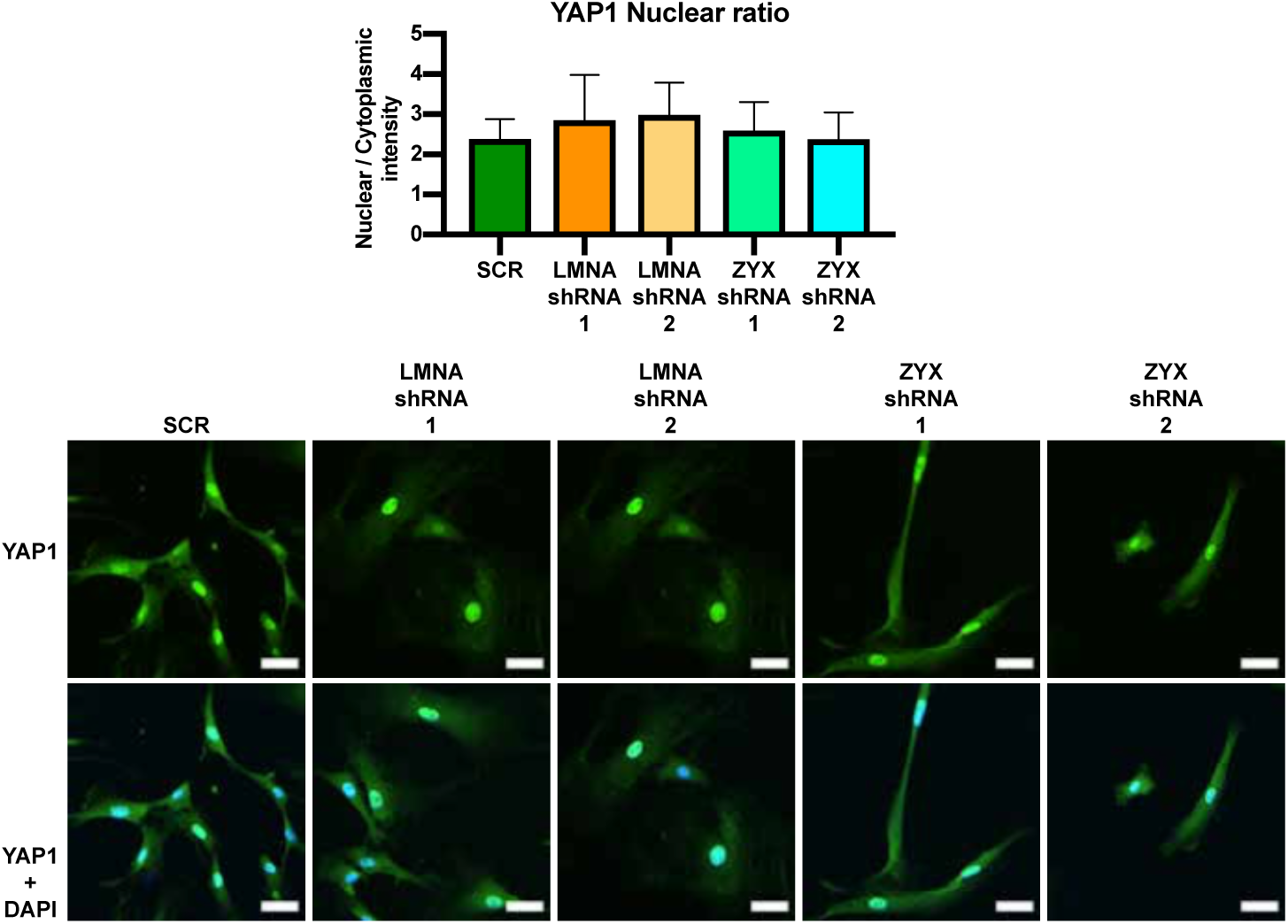
Lamin A and C and Zyxin do not regulate YAP1 nuclear localization. YAP1 staining (green) and nuclei (blue) of hMSCs transduced with scrambled-, LMNA- or ZYX-shRNA. Top panel shows YAP1 staining alone, while the bottom panel shows YAP1 and nuclei (blue). Scalebars 30µm. Graph shows the corresponding quantification of. Total cells analyzed for SCR: 20, LMNA-shRNA-1: 15, LMNA-shRNA-2: 21, ZYX-shRNA-1: 21, ZYX-shRNA-2: 17. One-way ANOVA, not significant. Error bars represent mean±SD.

